# Quantitative proteomic analysis of skeletal muscles from wild type and transgenic mice carrying recessive *Ryr1* mutations linked to congenital myopathies

**DOI:** 10.1101/2022.09.26.509474

**Authors:** Jan Eckhardt, Alexis Ruiz, Stéphane Koenig, Maud Frieden, Alexander Schmidt, Susan Treves, Francesco Zorzato

## Abstract

Skeletal muscle is a highly structured and differentiated tissue responsible for voluntary movement and metabolic regulation. Muscles however, are heterogeneous and depending on their location, speed of contraction, fatiguability and function, can be broadly subdivided into fast and slow twitch as well as subspecialized muscles, with each group expressing common as well as specific proteins. Congenital myopathies are a group of non-inflammatory non-dystrophic muscle diseases caused by mutations in a number of genes, leading to a weak muscle phenotype. In most cases specific muscles types are affected, with preferential involvement of fast twitch muscles as well as extraocular and facial muscles. Here we performed relative and absolute quantitative proteomic analysis of EDL, soleus and extraocular muscles from wild type and transgenic mice carrying compound heterozygous mutations in *Ryr1* identified in a patient with a severe congenital myopathy. Our quantitative proteomic study shows that recessive *Ryr1* mutations not only decrease the content of RyR1 protein in muscle, but also impact the content of many other proteins; in addition, we provide important insight into the pathological mechanism of congenital myopathies linked to mutations in other genes encoding components of the excitation contraction coupling molecular complex.

## INTRUCTION

Skeletal muscles constitute the largest organ, accounting for approximately 60% of the total body mass; they are responsible for movement and posture and additionally, play a fundamental role in regulating metabolism. Furthermore, skeletal muscles are plastic and can respond to physiological stimuli such as increased workload and exercise by undergoing hypertrophy. Broadly speaking muscles can be subdivided into different types depending on their speed of contraction, namely slow twitch muscles are characterized by level of oxidative activity, while fast twitch muscles show high content of enzymes involved in glycolytic activity. Fast- and slow-twitch muscle can be also identified based on the expression of specific myosin heavy chain (MyHC) isoforms (1, 2). Fast twitch muscles, also known as type II fibers, are specialized for rapid movements, are mainly glycolytic contain large glycogen stores and few mitochondria, fatigue rapidly and characteristically express the MyHC isoforms 2X, 2B and 2A. They are also the first muscles to appear during development and are more severely impacted in patients with congenital myopathies; they also undergo more prominent age-related atrophy or sarcopenia (1–6). Slow twitch muscles (type 1 fibers) are mainly oxidative, contain many mitochondria and are fatigue resistant. Slow twitch muscle, such as soleus, contain muscle fibers expressing the MyHC 1 isoform in addition of muscle fibers expressing MyHC 2A (2). Type 1 fibers are generally less severely affected in patients with neuromuscular disorders such congenital myopathies.

Although such a general classification based on MyHC isoform expression was used for many years by biochemists and physiologists, it has been recently improved thanks to the implementation of “omic” approaches which have helped refine the phenotypic signature at the single fiber level. A great deal of data has shown that type 2A fast fibers display a protein profile similar to type I fibers, namely a remarkable level of enzymes involved in oxidative metabolism. Interestingly, type 2X fibers apparently encode proteins annotated to both oxidative and glycolytic pathways (7, 8).

There are also a number of functionally specialized muscles including extraocular muscles (EOM), jaw muscles and inner ear muscles that have a different embryonic origin and are made up of atypical fiber types (2). For example, EOMs are the fastest contracting muscles yet they are fatigue resistant, contain many mitochondria and express most MyHC isoforms including type 1, embryonic and neonatal MyHC as well as EO-MyHC (9). EOMs are also specifically spared in patients with Duchenne Muscular Dystrophy yet they are affected in patients with some congenital myopathies, including patients with recessive *RYR1* myopathies carrying a hypomorphic or null allele (9–12).

Congenital Myopathies (CM) are a genetically heterogeneous group of early onset, non-dystrophic diseases preferentially affecting proximal and axial muscles. More than 20 genes have been implicated in CM, the most commonly affected being those encoding proteins involved in calcium homeostasis and excitation contraction coupling (ECC) and thin-thick filaments (13). Mutations in *RYR1*, the gene encoding the ryanodine receptor (RyR1) calcium channel of the sarcoplasmic reticulum, are found in approximately 30% of all CM patients, making it the most commonly mutated gene in human CM (12, 13). Within the group of patients carrying *RYR1* mutations, those with the recessive form of the disease are more severely affected, present from birth, have axial and proximal muscle weakness as well as involvement of facial and EOM (5, 12, 13). A common finding is also the reduced content of RyR1 protein in muscle biopsies (14, 15) which could be one of the causes leading to the weak muscle phenotype. To date, the pathomechanism of disease of recessive *RYR1* mutations is not completely understood and for this reason we created a mouse model knocked in for compound heterozygous mutations identified in a severely affected child. The double knock in mouse, henceforth referred to as double heterozygous or dHT mouse, carries the p.Q1970fsX16 mutation in one allele leading to the absence of a transcript due to nonsense-mediated decay of the allele carrying the frameshift mutation, and the mis-sense p.A4329D mutation in the other allele (16). The muscle phenotype of the dHT mouse model closely resembles that of human patients carrying a hypomorphic allele plus a mis-sense *RYR1* mutation, including reduced RyR1 protein content in skeletal muscles, the presence of cores and myofibrillar dis-array, mis-alignment of RyR1 and the dihydropyridine receptor and impaired EOM function (16, 17). Interestingly, beside a reduction in RyR1, the latter muscles also exhibited a significant decrease in mitochondrial number as well as changes in the expression and content of other proteins, including the almost complete absence of the EOM-specific MyHC isoform (17). Such results imply that broad changes in protein expression caused by the mutation and/or reduced content of RyR1 channels, impact other signaling pathways, leading to altered muscle function. A corollary to this is that since not all muscles are equally affected (for example fast twitch muscles and EOMs are more affected than slow twitch muscles) there may be differences in how the *RYR1* mutations affect the different muscle types.

In order to establish how and if *Ryr1* mutations differentially impinge on the expression and function of proteins specific for different muscle types, we performed qualitative and quantitative proteomic analysis of EDL, soleus and EOMs from wild type and dHT mice.

## RESULTS

Figure 1 shows a diagram of our experimental work flow: three muscle types were isolated from 12 weeks old wild type (n=5) and dHT (n=5) mice, samples were processed for Mass Spectrometry and the results obtained were analyzed against a protein database containing sequences of the predicted SwissProt entries of *mus musculus* (www.ebi.ac.uk, release date 2019/03/27), Myh2 and Myh13 from Trembl, the six calibration mix proteins (18) and commonly observed contaminants (in total 17,414 sequences) using the SpectroMine software. Results obtained from five muscles per group were averaged, filtered so that only changes in protein content greater than 0.25-fold and showing a significance of q<0.05 or greater, were considered. In addition, proteins yielding only 1 peptide were not used for analysis and were filtered out.

**Figure 1:**
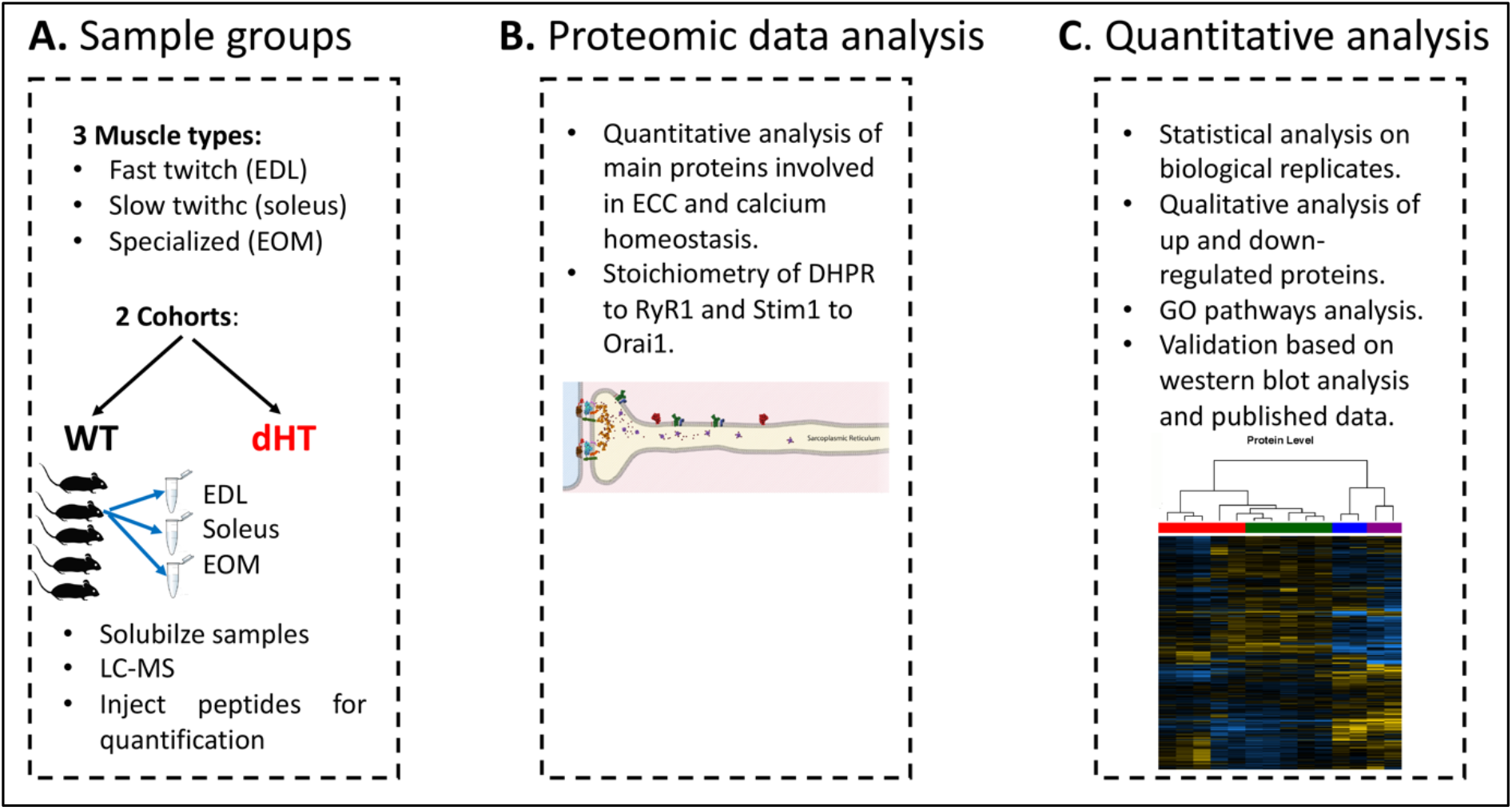
Schematic overview of the workflow. **A.** Skeletal muscles from 12 weeks old WT (5 mice) and dHT littermates (5 mice) were isolated and flash frozen. Three different types of muscles were isolated per mouse, namely EDL, soleus and EOMs. On the day of the experiment, muscles were solubilized and processed for LC-MS. **B.** For absolute protein quantification, synthetic peptides of RyR1, Cav1.1, Stim1 and Orai1 were used. **C**. Protein content in different muscle types and in the different mouse genotypes were analyzed and compared.

### Comparison of the proteome of EDL and soleus muscles from WT mice

In order to perform their specific physiological functions, different muscle types express different protein isoforms or different amounts of specific proteins. For example, slow twitch muscles contain large amounts of the oxygen binding protein myoglobin and of carbonic anhydrase III the enzyme catalyzing the conversion of CO_2_ to H_2_CO_3_ and HCO_3_^-^ (19, 20), while fast twitch muscles express large amounts of the calcium buffer protein parvalbumin (21); additionally, each muscle type contains specific isoforms of contractile and sarcomeric proteins (2). Our first aim was to analyze the proteomes of wild type mouse EDL and soleus muscles to establish their most important qualitative differences.

Figure 2A shows that the content of more than 1800 proteins varies significantly (q<0.05) between EDL and soleus muscles from WT mice, of these 547 are present in lower amounts and 1319 are present in higher amounts in soleus compared to EDL muscles; figure 2B shows a volcano plot of the log_2_ fold change of proteins in slow (condition 2) versus fast (condition 1) muscles. Enriched GO pathway analysis of the molecular function pathways revealed that there is a significant decrease of “oxidoreductase activity associated genes” in fast twitch fibers, a not unexpected finding considering that slow twitch muscles are made up type I and type IIa/IIx fibers which contain more mitochondria and oxidative enzymes than fast twitch type IIb fibers of fast twitch muscles. To have a broader view of the overall differences in the two muscle types we analyzed the GO “Reactome pathway” terms (Fig.2C); in this case, changes in additional pathways were observed including (i) a decrease in EDL muscles of pathways associated with mitochondrial function (fatty acid metabolism, TCA cycle, electron transport chain, complex 1 biogenesis and ß-oxidation); (ii) changes in proteins involved in muscle contraction in both EDL and soleus muscles, (iii) an increase in EDL muscles of collagen, integrins and extracellular matrix proteins.

**Figure 2:**
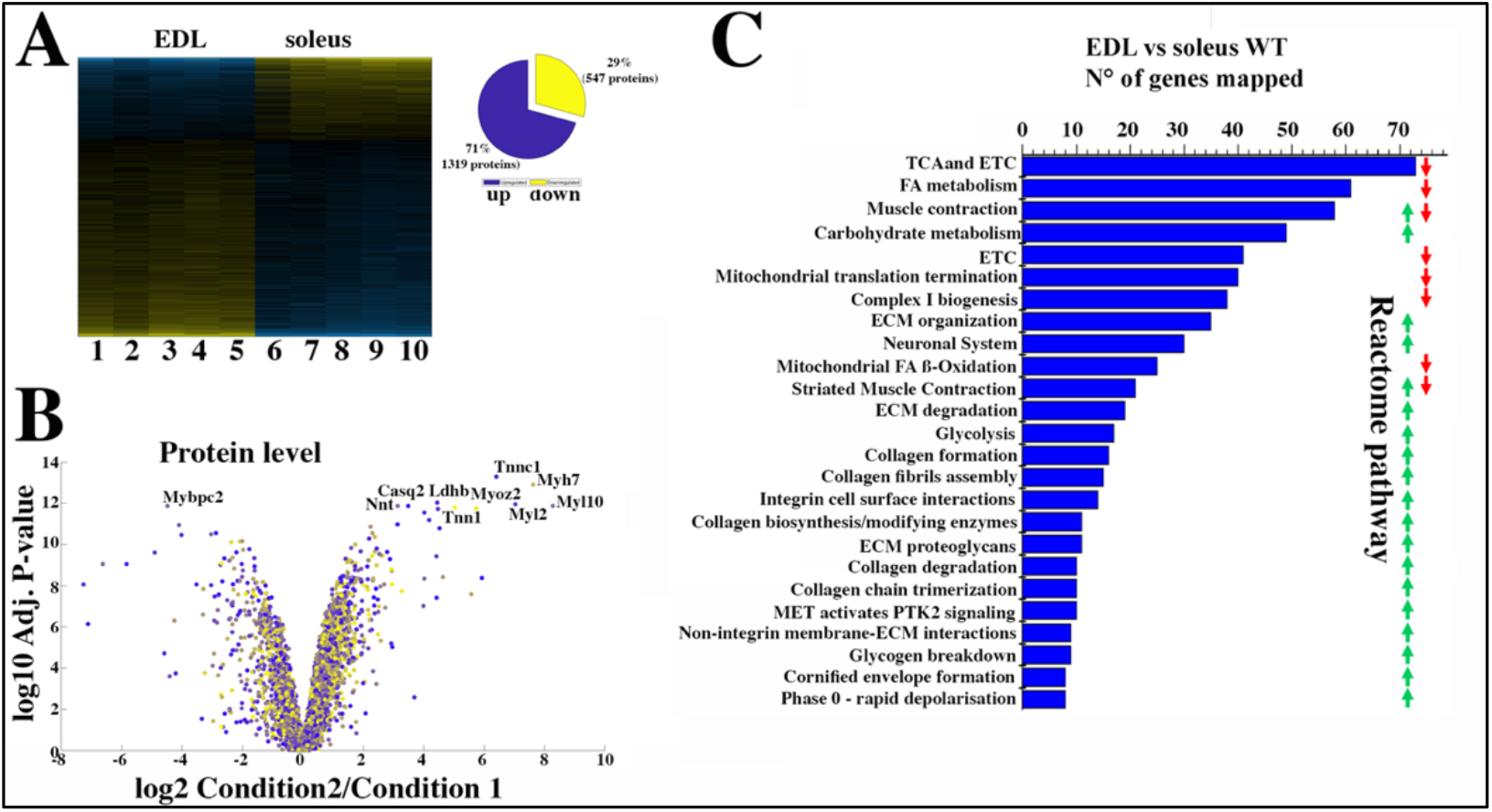
Proteomic analysis of EDL and soleus muscles from WT mice confirms the significant difference in content if proteins involved in the TCA cycle and electron transport chain, fatty acid metabolism and muscle contraction. **A.** Hierarchically clustered heatmaps of the relative abundance of proteins in EDL and soleus muscles from 5 mice. Blue blocks represent proteins which are increased in content, yellow blocks proteins which are decreased in content in EDL versus soleus muscles. Right pie chart shows overall number of increased (blue) and decreased (yellow) proteins. Areas are relative to their numbers. **B.** Volcano plot of a total of 1866 quantified proteins which showed significant increased (blue) and decreased (yellow) content. The horizontal coordinate is the difference multiple (logarithmic transformation at the base of 2), and the vertical coordinate is the significant difference p value (logarithmic transformation at the base of 10). The proteins showing major change in content are abbreviated. **Soleus: condition 2**; **EDL: condition 1 C.** Reactome pathway analysis showing major pathways which differ between EDL and soleus muscles. A q-value of equal or less than 0.05 was used to filter significant changes prior to the pathway analyses.

However, Genome Ontology pathway analysis is not sufficiently informative and probably misses important groups of proteins specific to skeletal muscle function; this observation prompted us to select specific proteins whose expression level is known to be different between fast and slow twitch muscles. Focusing on the content of contractile and sarcomeric proteins our results confirm that the slow muscle Troponin I and C1 isoforms as well as MyHC 1 (encoded by Myh7) are enriched between 32 and 197-fold in soleus muscles, whereas a-actinin 3 and 4 and myomesin 1 are more abundant in EDL muscles and desmin is enriched in soleus muscles (Table 1). Analysis of sarcoplasmic reticulum proteins involved in ECC showed that the content of calsequestrin 2 and SERCA2 is 11- and 22-fold higher in soleus muscles, whereas the relative content of the RyR1, the dihydropyridine (DHPR) complex (including the α1, β1 and α2δ subunits), Stac3 and triadin is more than 50% higher in EDL muscles compared to soleus, as is FKBP12 which binds to and stabilizes the RyR1 complex (22). Fast twitch muscles are also enriched in SERCA1, calsequestrin 1 and junctophilin 1 and 2. Interestingly, EDL are also enriched in proteins annotated to “calcium dependent signaling” via the calcium /calmodulin dependent protein kinase IIa and IIg. On the other hand, more than 10 heat shock proteins are more abundant in soleus muscles, including Hsp70. The latter protein has been implicated in expression of Glut4 in slow twitch muscles (23). Importantly, a great deal of data have shown that muscles from patients with several neuromuscular disorders including those caused by *RYR1* mutations show fiber type 1 predominance (4, 5) and Hsp70 has been reported to be involved in a variety of mechanism enhancing cell survival (24).

**Table 1:**
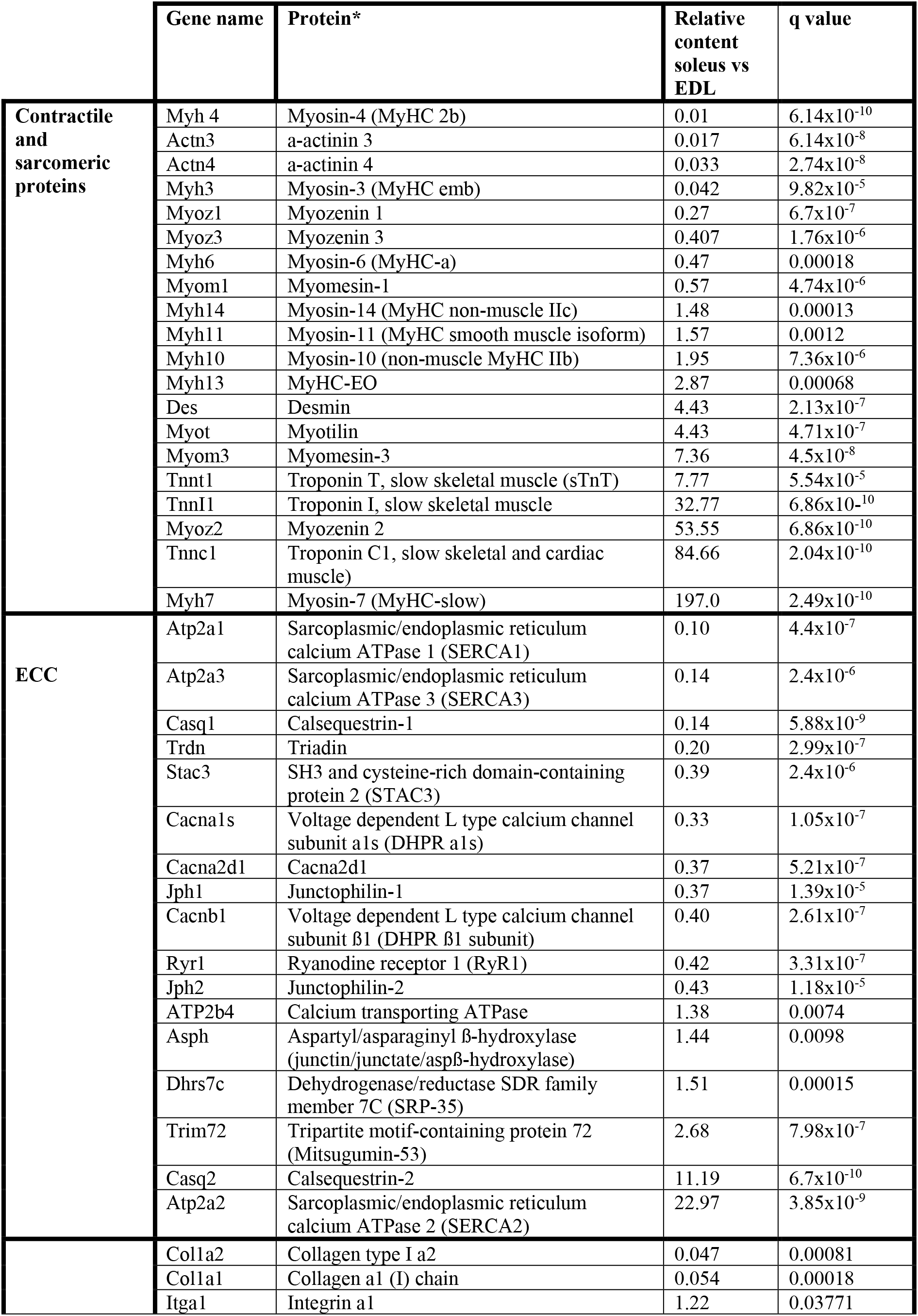
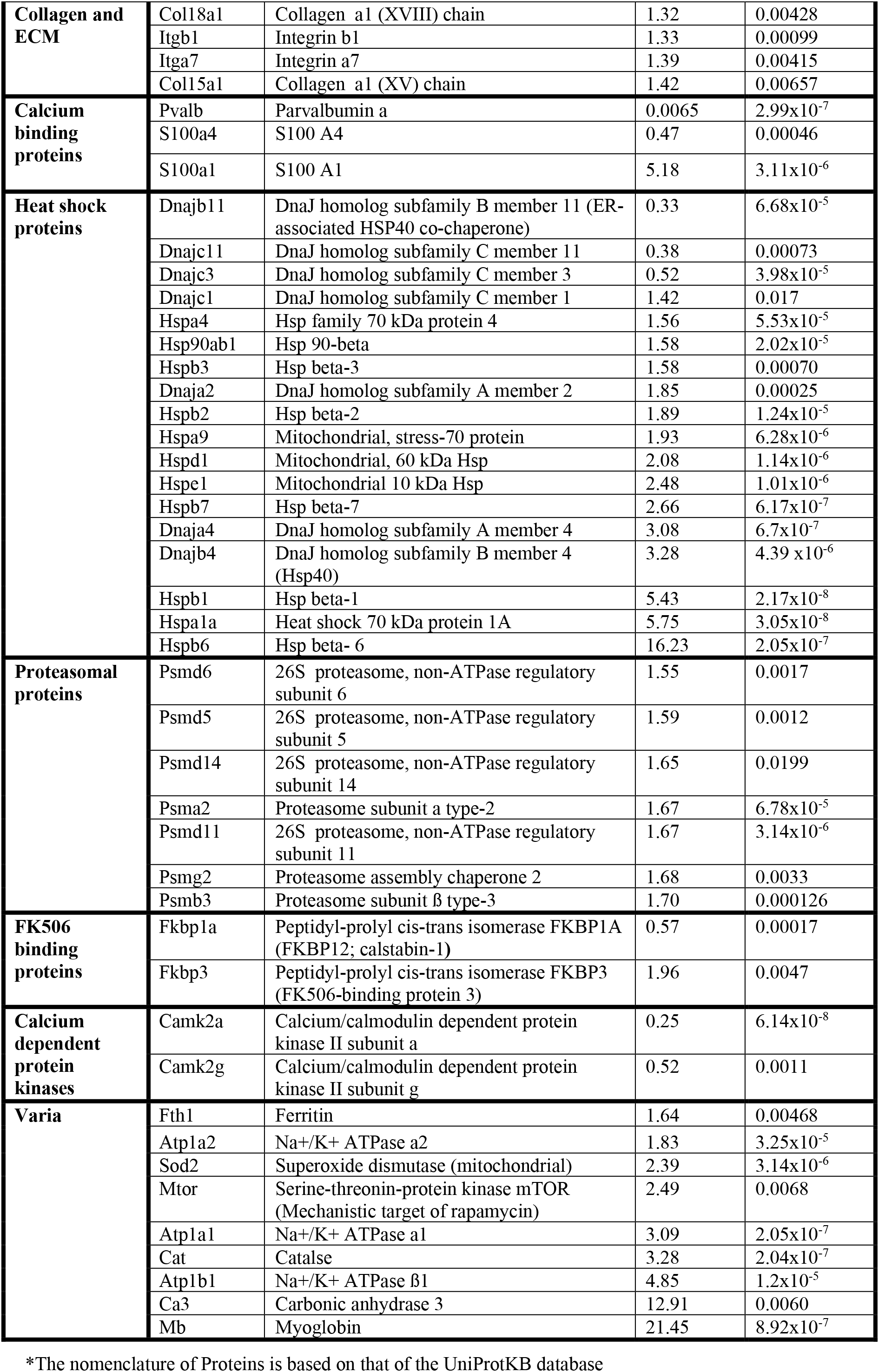
Relative change in protein content between soleus and EDL (baseline) muscles isolated from WT mice.

Furthermore, the content of mitsugumin 53 (encoded by *Trim72*), a protein involved in muscle membrane repair (25) is 2.8 fold higher in slow twitch muscles compared to fast twitch muscles. Thus, on the basis these observations we cannot exclude the possibility that increased expression of Hsp70 and/or of other proteins such as mitsugumin 53 might be relevant in preventing muscle fiber type 1 damage associated with the presence of recessive *RYR1* mutations or with other type of stressing events (26). To verify this hypothesis, we next examined the proteome of fast and slow twitch muscles in a mouse model (RyR1 dHT) for neuromuscular disorders carrying the p.Q1970fsX16 mutation in one allele and the mis-sense p.A4329D mutation in the other allele (16).

### Comparison of muscles isolated from WT and RyR1 dHT mice

In the next experiments the proteome of three different muscles from WT mice vs those of dHT mice were compared. Fig. 3A and B shows that in EDL muscles a total of 848 proteins are significantly (q<0.05) mis-regulated in dHT mice; in particular, 529 and 319 proteins are up- or downregulated only in the EDLs of dHT mice compared to WT mice, respectively (Supplementary Fig. 2). GO pathway analysis revealed that proteins involved in homeostasis of the extracellular matrix, including collagen assembly and chain formation, collagen degradation, ECM organization and integrin interaction, are down-regulated in EDLs from the dHT mice. Since GO analysis appears to be not informative concerning changes occurring in pathways important for skeletal muscle function, we again selected and analyzed protein families playing a role in skeletal muscle ECC, muscle contraction, collagen and ECM, heat shock response/chaperones, protein synthesis and calcium-dependent regulatory functions. Table 2 shows that several proteins involved in skeletal muscle ECC are down-regulated, including the RyR1 as well as its stabilizing binding protein FKBP12, the α1 and ß1 subunits of the DHPR and junctophillin 1 whose relative content decreases by 30%, 23% and 40%, respectively. Asph which encodes different proteins including junctin, junctate, humbug and aspartyl-ß-hydroxylase (27) increases almost 2-fold, whereas calsequestrin 1 and SRP-35 (Dhrs7c) increase by 20 and 34% in EDLs from dHT mice. Additionally, the expression of type 2 fibers is impacted since MyHC 2X and 2B as well as α-actinin 3 (which is preferentially expressed in type 2 fibers) (2) are decreased in the EDLs of dHT mice. The decrease of the fast isoforms of MyHC in dHT EDL muscles is accompanied by a decrease of many collagen isoforms. On the other hand, the content of several heat shock proteins as well as the content of 60S and 40S ribosomal proteins is increased in fast twitch fibers from the dHT. In addition, we found that the calcium/CaM dependent protein kinases 1, 2α 2β and 2δ are increases in EDL from dHT mice.

**Figure 3:**
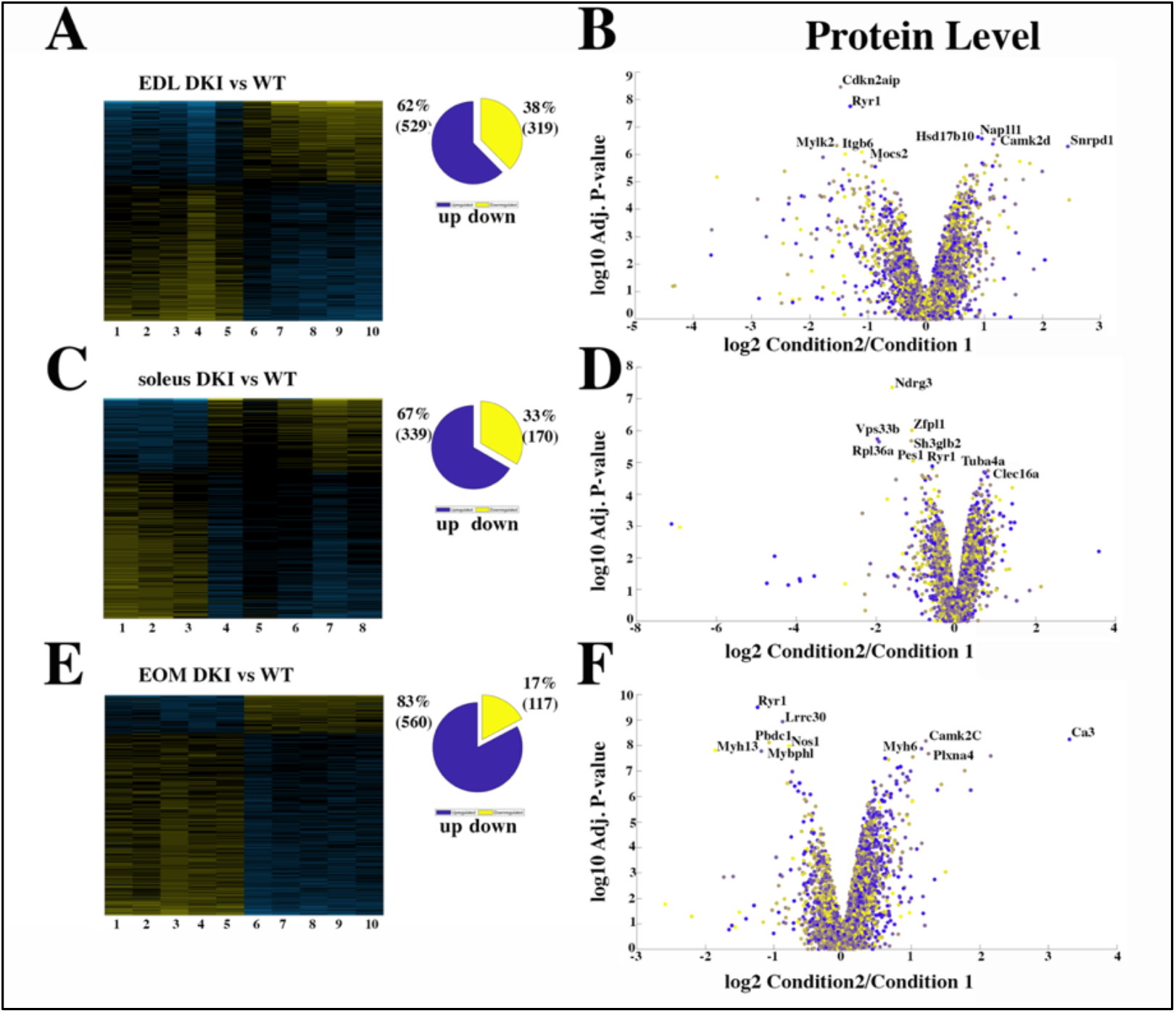
Proteomic analysis comparison of muscles from WT and dHT mice. **A, C and E.** Hierarchically clustered heatmaps of the relative abundance of proteins in EDL (A), soleus muscles (C) and EOMs (E) from 5 mice. Blue blocks represent proteins which are increased in content, yellow blocks proteins which are decreased in content in dHT versus WT. Right pie chart shows overall number of increased (purple) and decreased (yellow) proteins. Areas are relative to their numbers. **B, D and F** Volcano plots of total quantified proteins showing significant increased (blue) and decreased (yellow) content in dHT (condition 2) versus WT (condition 1) EDL (B), soleus (D) and EOMs (F). The horizontal coordinate is the difference multiple (logarithmic transformation at the base of 2), and the vertical coordinate is the significant difference p value (logarithmic transformation at the base of 10). The proteins showing major change in content are abbreviated.

**Table 2:**
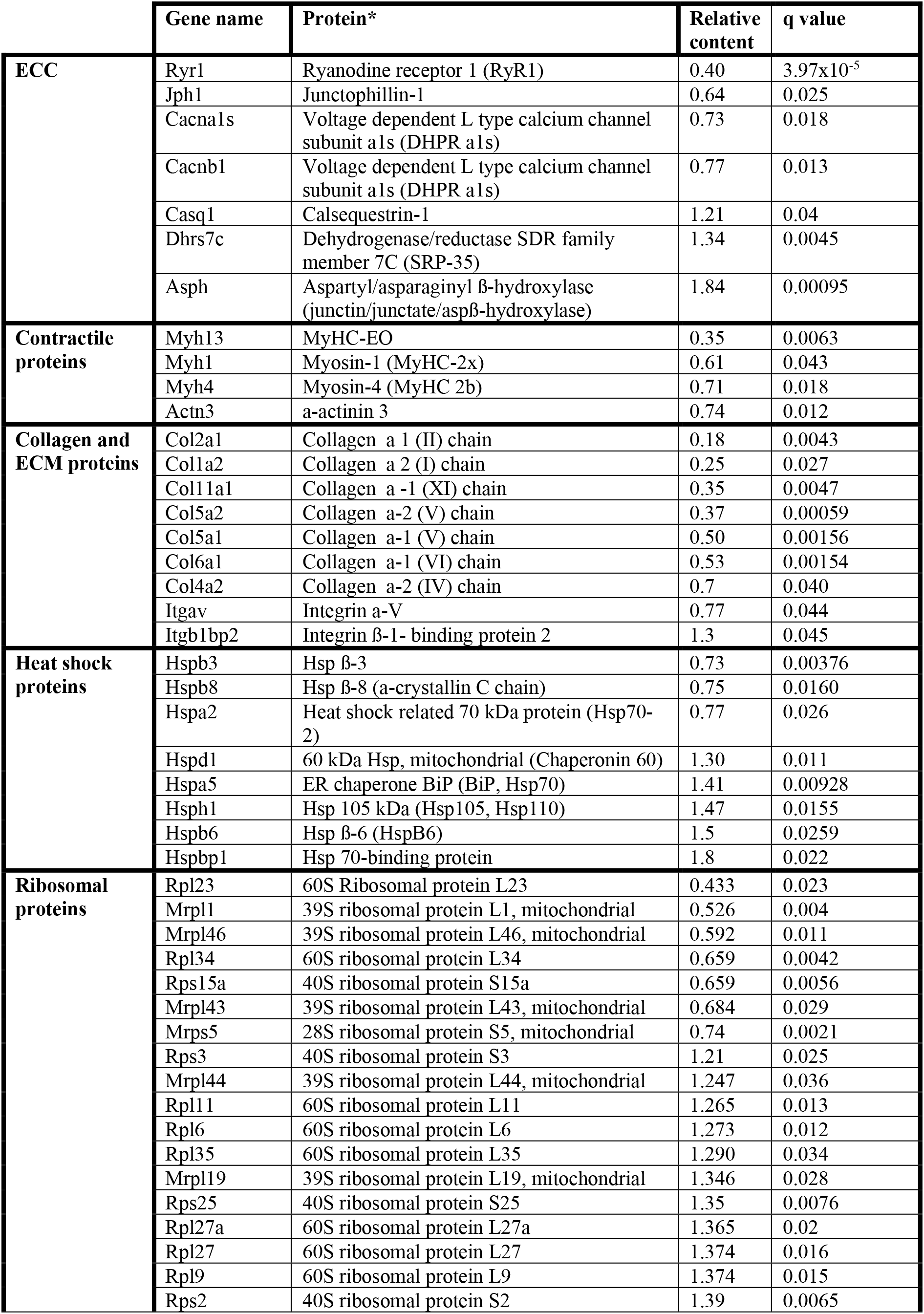
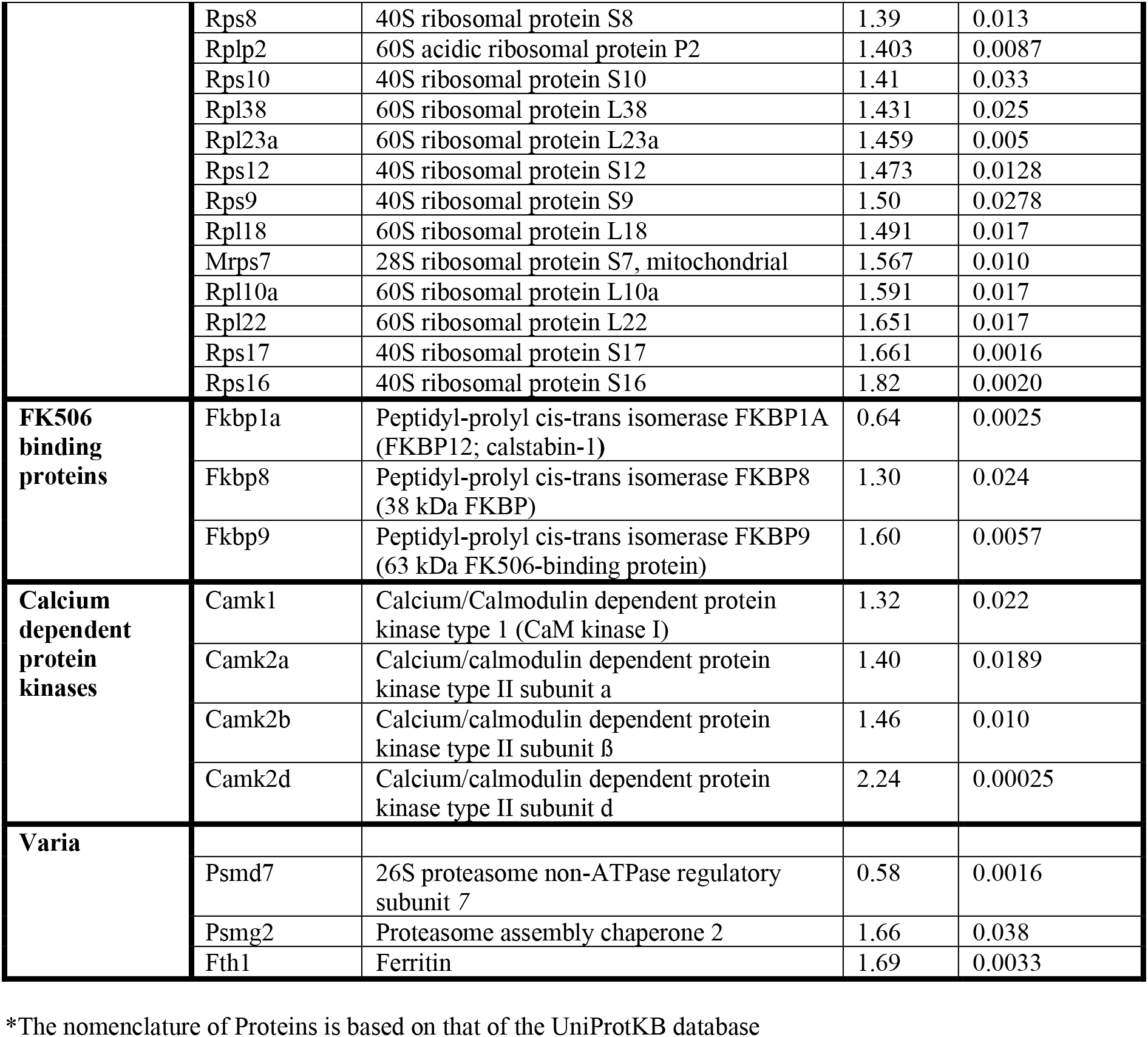
Relative change in the content of selected proteins in EDL muscles isolated from WT (baseline) and dHT mice.

We next compared the proteome of soleus muscles from WT and dHT mice. Fig.3C and D show that the overall number of proteins showing significant changes in their relative content between WT and dHT mice, is smaller than that observed in EDL muscles. In particular, we found that 339 and 170 proteins are up- or downregulated only in the soleus muscles of dHT mice compared to WT mice, respectively (Supplementary Fig 2). Contrary to EDL muscles, GO analysis failed to identify a preferentially affected cellular pathway so, as described in the previous sections, we selected and analyzed specific protein families that showed significantly different (q<0.05) content between the two mouse genotypes (Table 3). In the ECC protein category RyR1, Cav1.1 and Junctophillin 1 are significantly decreased, as is triadin, whereas junctin/junctate/ß-hydroxylase and SERCA2 are increased. In the contractile protein group, significant changes are only observed for Troponin 3 whose content decreases by about 30%. Similar to what was observed in EDL muscles, we found that the content of calcium/calmodulin dependent protein kinases IIδ and γ is increased. In addition, S100A1, a calcium binding protein which binds to and regulates RyR1 activity (28, 29), is significantly increased in soleus muscles from dHT mice. Finally, proteins constituting the 60S and 40S ribosomal subunits are increased in soleus muscles from dHT mice compared to WT.

**Table 3:**
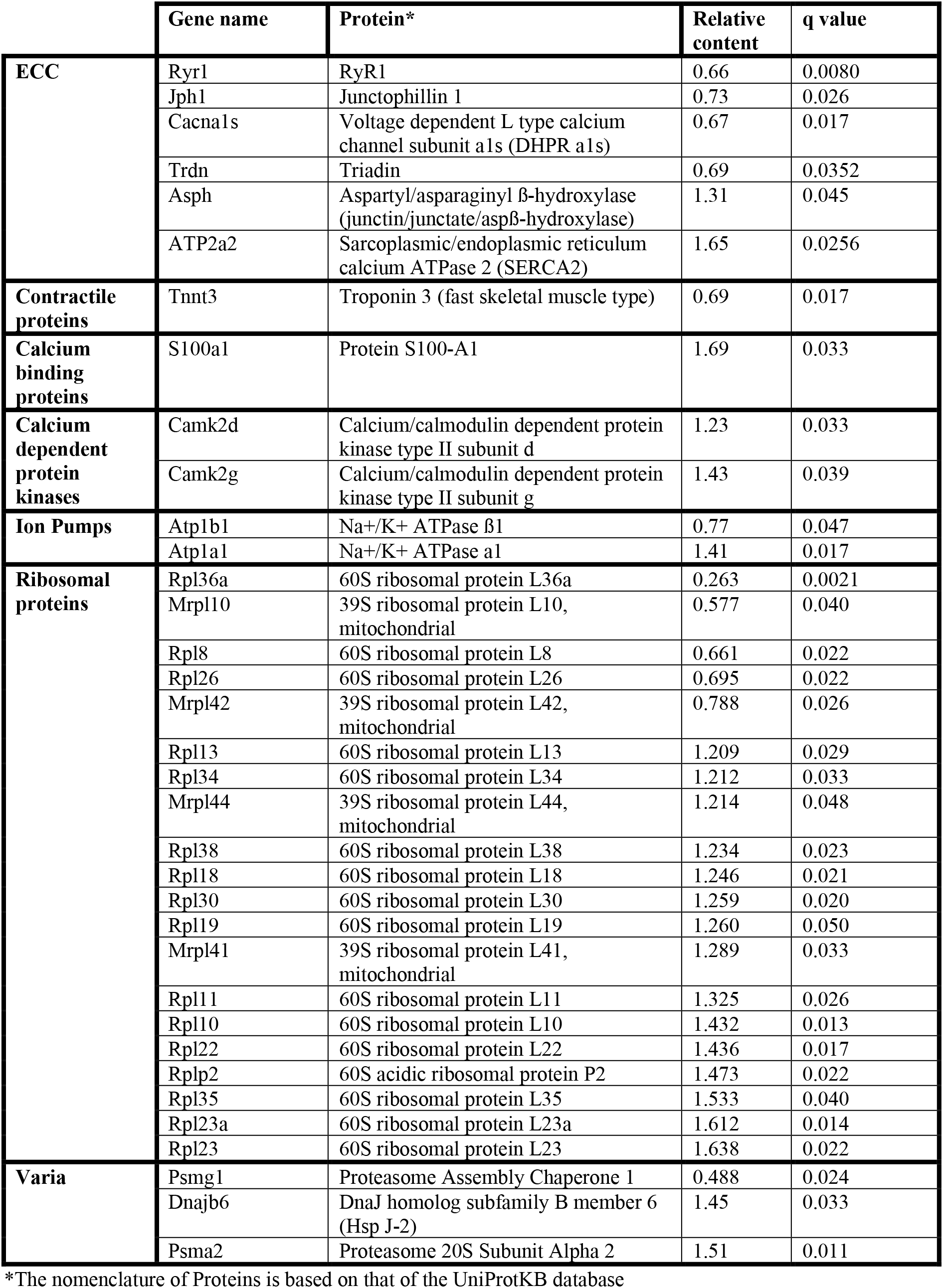
Relative change in the content of selected proteins in soleus muscles isolated from WT (baseline) and dHT mice.

Since ophthalmoplegia is a common clinical sign observed in patients affect by congenital myopathies linked to recessive *RYR1* mutations (5, 12, 13), we also investigated the proteome of EOMs from WT and dHT mice. Fig. 3E and F shows that 560 and 117 proteins are up- or downregulated only in the EOM of mutant dHT mice compared to WT mice, respectively (supplementary Fig 2). Interestingly, the overall changes caused by *Ryr1* mutations on the protein composition of muscles is more prominent in fast twitch muscles such as EOMs and EDLs compared to the slow twitch soleus muscle. In EOMs, the proteins showing the greatest fold change (aside those involved in ECC and muscle contraction), are heat shock proteins, ribosomal proteins and proteins of the ECM and collagen. Interestingly, in EOM muscles the content of proteins belonging to the collagen family are significantly increased in dHT versus WT, whereas they are decreased in the EDLs of dHT versus WT mice. In agreement with data obtained in EDL and soleus muscles, a variety of heat shock proteins and calcium/calmodulin dependent protein kinases IIβ and IIδ and S100 family proteins are more abundant in EOM form dHT versus WT mice.

The above analysis revealed that the content of many proteins differs between WT and dHT EDL, soleus and EOM muscles. We next refined our analysis and searched for protein whose content variation is most strongly associated with the dHT genotype. In particular, we searched for proteins which show significant changes in content in all three muscle types, namely EDL, Sol and EOM. The Venn diagram (Supplementary Fig.2) shows that the three muscle types from the dHT mice share a number of proteins whose content increases or decreases. The downregulation of RyR1 appears to be a unique a signature of the dHT phenotype, since its decrease is the only change shared between EDL, soleus and EOM (Supplementary Fig. 2). Furthermore, other proteins annotated to calcium signaling such as calmodulin kinase 2δ and aspartyl-beta-hydroxylase are increased in content in all three muscle types from dHT mice, as are several proteins associated with the 40S and 60S ribosomal subunits. Two heat shock proteins Hsp70 (BiP) and Hsp family B small member 6 (HSPB6) are increased only in EDL and EOMs.

### Quantification and stoichiometry of ECC proteins in WT and dHT muscles

Skeletal muscle ECC relies on the highly ordered architecture of two intracellular membrane compartments, namely the transverse tubules which are invaginations of the plasma membrane containing the DHPR macromolecular complex and the sarcoplasmic reticulum containing the RyR1 macromolecular complex, as well as other proteins involved in calcium homeostasis and accessory structural proteins (30, 31). The relative content of many of these proteins has been determined, nevertheless few studies have established their stoichiometry in relation to particular muscle types (31–33). Within a total muscle homogenate, sarcoplasmic reticulum membrane proteins are of low abundance, thus, to quantify these proteins we performed high resolution TMT mass spectrometry by using spiked-in labeled peptides from major protein involved in key steps of ECC calcium signaling to build a standard calibration curve. In particular, we used peptides from RyR1 and Cacna1s, and Stim1 and Orai1, proteins which are involved in calcium release from the SR and in calcium entry across sarcolemma, respectively. The obtained protein concentrations showed a high correlation (R2=0.96, Supplementary Fig. 3) with the MS abundance estimates determined from the global proteomics analysis. Therefore, we used this curve to extrapolate the absolute amounts and stoichiometry of proteins whose values fall within the linear domain of the curve, namely, JP-45, triadin, junctophilin 1, Stac3 in addition to RyR1, Cacna1s, Stim1 and Orai1. The content of the RyR1 protomer in WT fast twitch EDL muscles is 1.29±0.07 μmol/kg wet weight, whereby the calculated RyR1 tetrameric complex is 0.32 μmol/kg wet weight (Table 5) a value which is 3-fold lower compared to that determined in total muscle homogenates by[^3^H]-ryanodine equilibrium binding by Bers et al. (34). On the other hand, our RyR1 quantification results in mouse total muscle homogenates obtained by TMT mass spectrometry using labeled peptides is approximately 5-fold higher compared to those obtained in rabbit and frog whole skeletal muscle homogenate preparations by Anderson et al. (35) and by Margreth et al. (36). Our results also show that the RyR1 concentration (in μmol/Kg) in soleus and EOM muscles from WT mice is approximately 38% and 46% of that found in EDL muscles of WT mice, respectively (Table 5). We found that the content of Cacna1s in EDL muscles is 0.56±0.03 μmol/kg wet weight, a value approx. 2.5-fold higher compared to that of soleus muscles (0.18±0.01 μmol/kg wet weight) and of EOMs (0.21± 0.01 μmol/kg wet weight). Thus, the calculated RyR1 tetramer to Cacna1s ratio in EDL muscles from WT and dHT mice is 0.571 and 0.429, respectively (Table 6). Such a value appears to be slightly higher both in soleus and EOM muscles (0.667 and 0.625 in WT and dHT soleus muscles and 0.714 and 0.474 in WT and dHT EOMs, respectively, Table 6). The Stac3 content correlates with that of Cacna1s, namely EDL is the muscle which is most enriched in Stac3 (0.62±0.07 μmol/kg wet weight); soleus and EOMs contain approximately one third of the Stac 3 present in the EDL, namely 0.22±0.02 μmol/kg wet weight and 0.17±0.01 μmol/kg wet weight in soleus and EOMs, respectively. Stac3 content in muscles from dHT was similar to that of WT littermates.

**Table 4:**
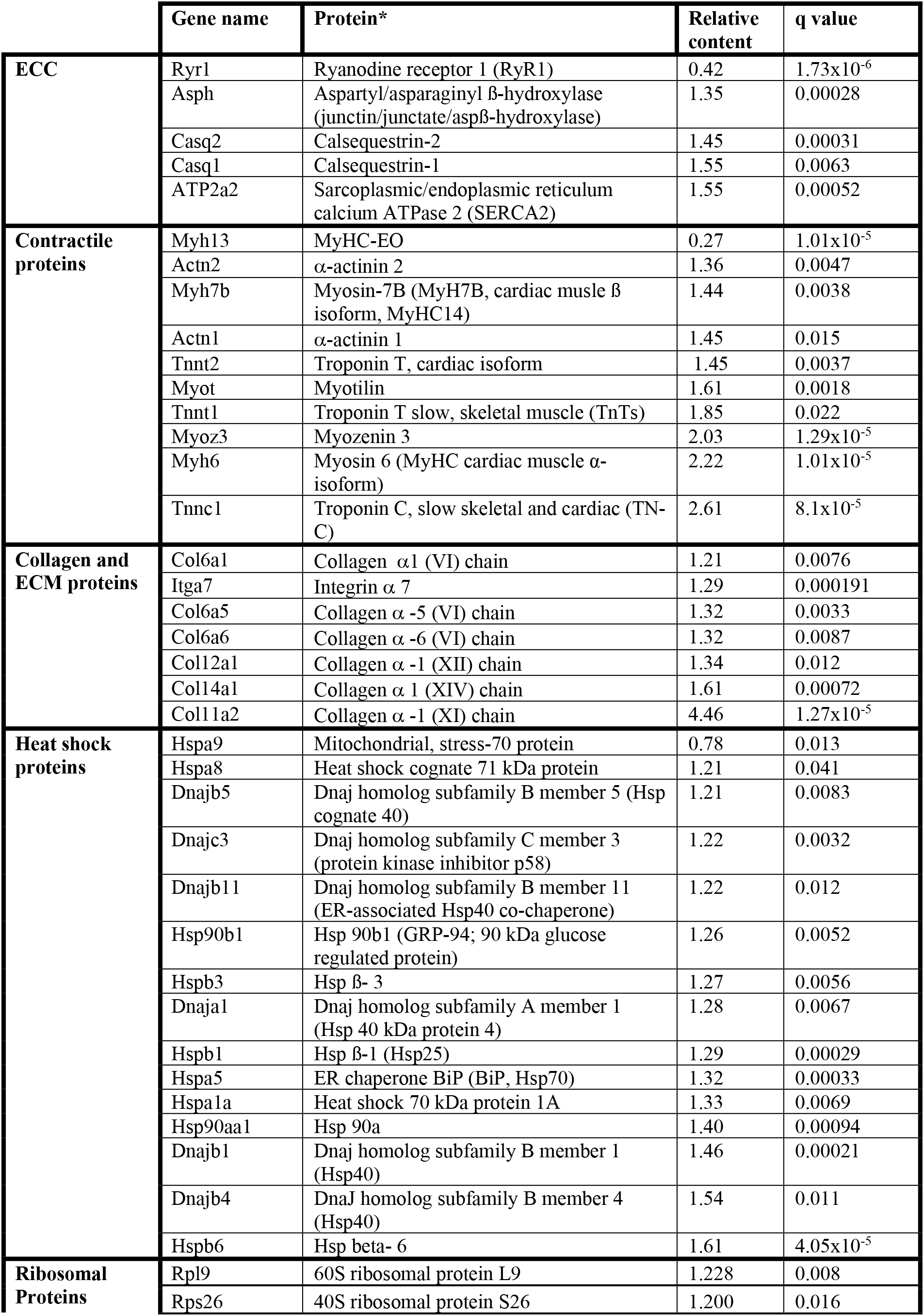
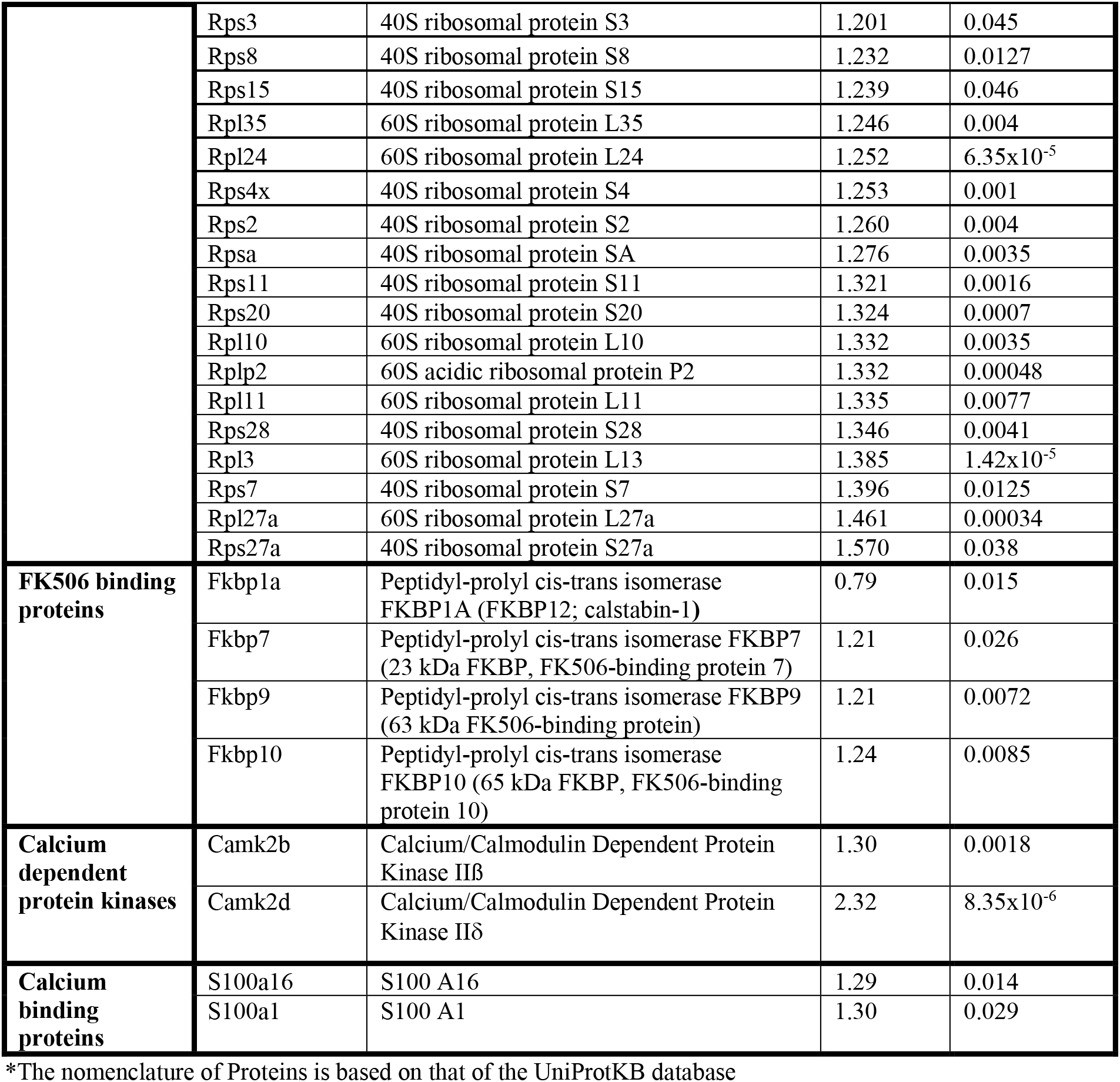
Relative change in the content of selected proteins in EOM isolated from WT and dHT mice.

**Table 5:**
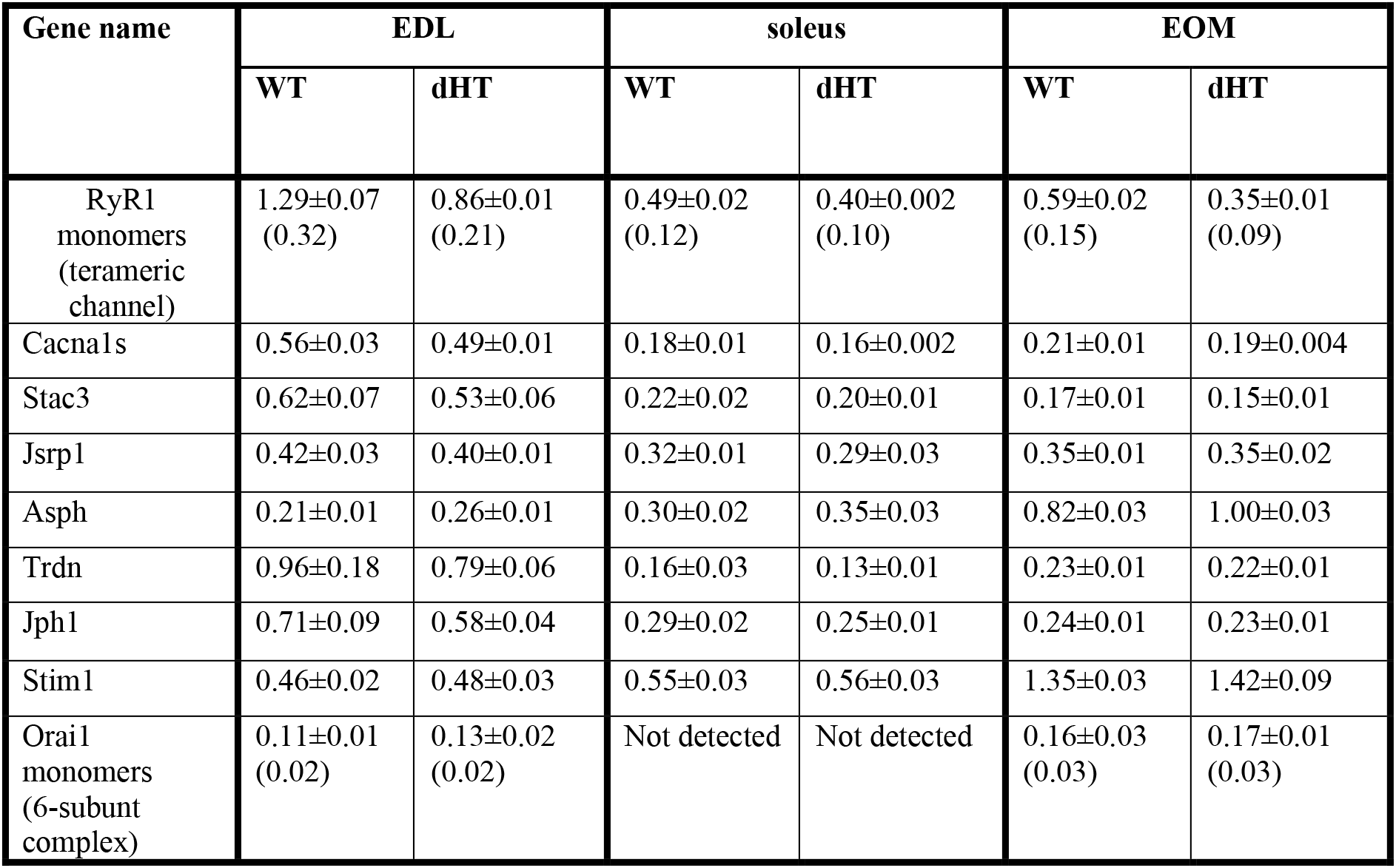
Concentration μmol/Kg (mean±SD) of proteins involved in ECC in EDL, soleus and EOM muscles from WT (n=5 mice) and dHT (n=5 mice) using the peptide 4 point calibration curve.

| Gene name | EDL |  | soleus |  | EOM |  |
| --- | --- | --- | --- | --- | --- | --- |
|  | WT | dHT | WT | dHT | WT | dHT |
| RyR1 monomers (terameric channel) | 1.29±0.07 (0.32) | 0.86±0.01 (0.21) | 0.49±0.02 (0.12) | 0.40±0.002 (0.10) | 0.59±0.02 (0.15) | 0.35±0.01 (0.09) |
| Cacna1s | 0.56±0.03 | 0.49±0.01 | 0.18±0.01 | 0.16±0.002 | 0.21±0.01 | 0.19±0.004 |
| Stac3 | 0.62±0.07 | 0.53±0.06 | 0.22±0.02 | 0.20±0.01 | 0.17±0.01 | 0.15±0.01 |
| Jsrp1 | 0.42±0.03 | 0.40±0.01 | 0.32±0.01 | 0.29±0.03 | 0.35±0.01 | 0.35±0.02 |
| Asph | 0.21±0.01 | 0.26±0.01 | 0.30±0.02 | 0.35±0.03 | 0.82±0.03 | 1.00±0.03 |
| Trdn | 0.96±0.18 | 0.79±0.06 | 0.16±0.03 | 0.13±0.01 | 0.23±0.01 | 0.22±0.01 |
| Jph1 | 0.71±0.09 | 0.58±0.04 | 0.29±0.02 | 0.25±0.01 | 0.24±0.01 | 0.23±0.01 |
| Stim1 | 0.46±0.02 | 0.48±0.03 | 0.55±0.03 | 0.56±0.03 | 1.35±0.03 | 1.42±0.09 |
| Orai1 monomers (6-subunit complex) | 0.11±0.01 (0.02) | 0.13±0.02 (0.02) | Not detected | Not detected | 0.16±0.03 (0.03) | 0.17±0.01 (0.03) |

**Table 6:**
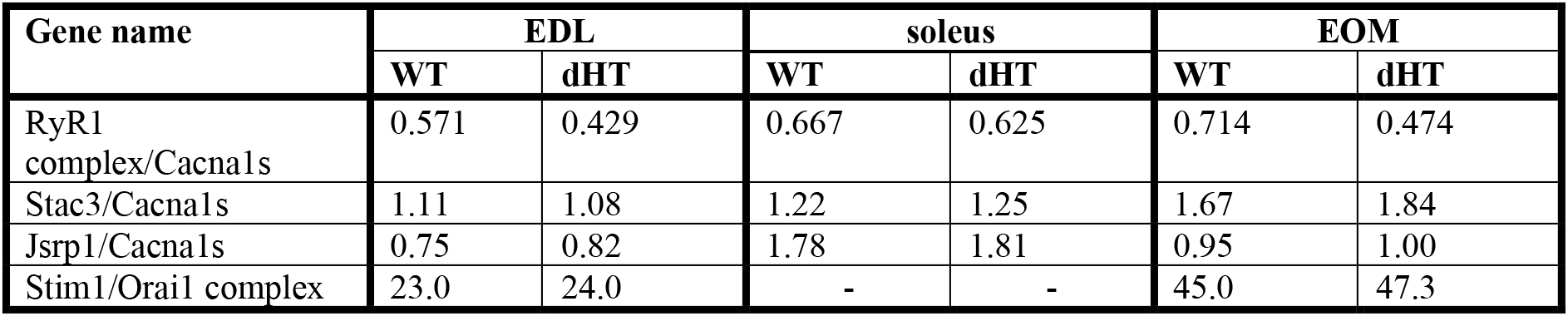
calculated ratio values

Interestingly, the content of Stim1 depends on the muscles type. Mass spec quantification revealed that EOMs contain the highest amounts of Stim1 (1.35±0.03 μmol/kg wet weight) compared to soleus (0.55±0.03 μmol/kg wet weight) and EDL (0.46±0.02 μmol/kg wet weight). Western blot analysis of total muscle homogenates from WT mice confirmed that EOMs contain 4 times more Stim1 than EDL muscles (Fig. 4) and that equal proportions of Stim1 and Stim1L are present in the three muscle groups, with no preferential expression of the long isoform in any of the muscles investigated. As to WT EDL and soleus, we found no major differences in Stim1 expression, confirming previous data by Cully et al. (37). The expression of Stim1 is accompanied by the expression of Orai1 in EDL and EOMs but not in soleus muscles. Indeed, mouse EOMs contain the highest amount of Orai1 monomer (0.16±0.03 μmol/kg wet weight) and EDLs contained approximately 68% of that (0.11±0.01 μmol/kg wet weight). To our surprise in soleus muscles, the content of Orai1 is below the detection level of mass spectrometry measurement, indicating that slow twitch (soleus) muscles express very little, if any, Orai1 compared to fast twitch EDL and EOMs.

**Figure 4:**
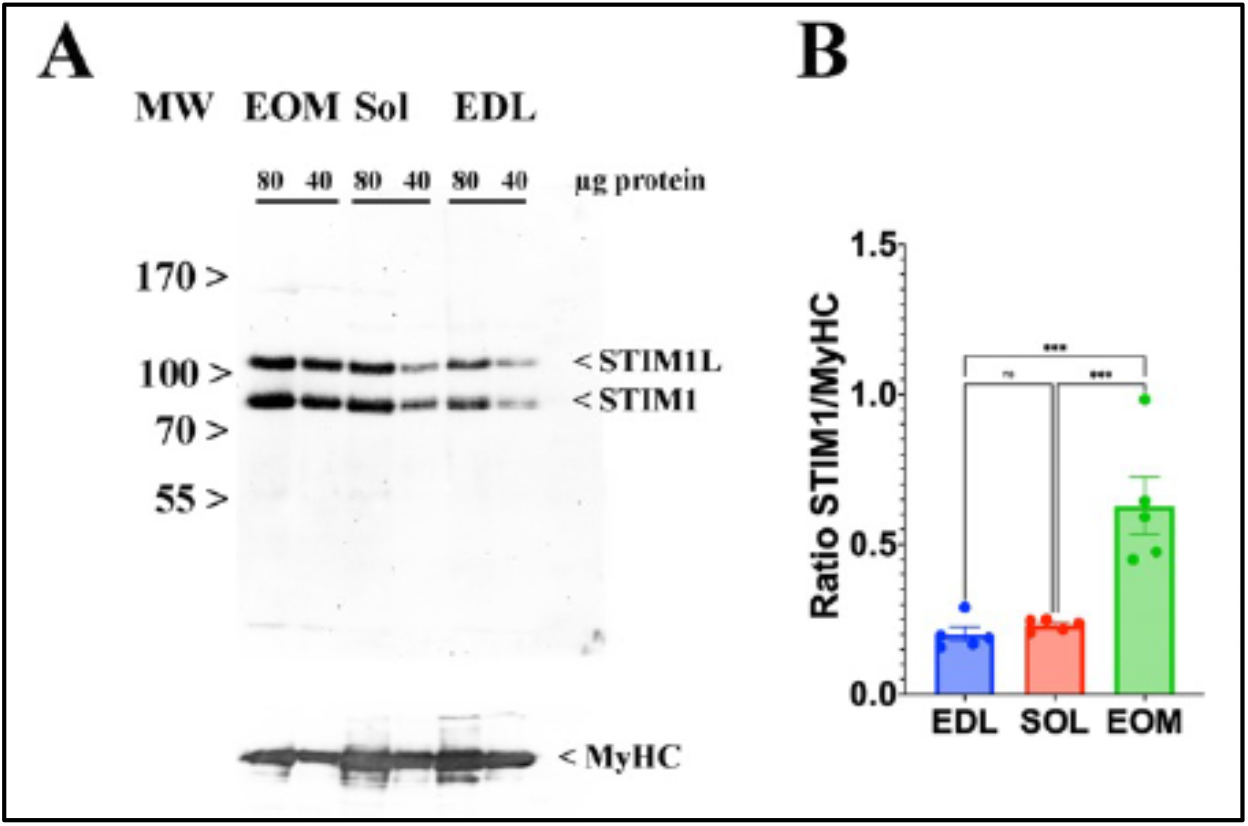
EOMs are enriched in Stim 1. **A.** Representative western blots showing Stim1 and Stim1L immunopositive bands. Forty and eighty micrograms of total homogenates from EOM, soleus, and EDL muscles isolated from WT mice were loaded onto a 7.5% SDS PAGE. Proteins were blotted onto nitrocellulose, probed with an antibody recognizing Stim1 and Stim1L, followed by incubation with an anti-rabbit IgG HRP-linked antibody. Bands were visualized by chemiluminescence. Blots were subsequently stripped and probed with anti-MyHC (all) for loading normalization (bottom panel). **B**. Relative content of Stim1 in the three muscle types examined. Each symbol represents the value of a single mouse. *** P<0.001.

## DISCUSSION

To understand in greater detail the changes in skeletal muscle function in congenital myopathies caused by recessive *RYR1* mutations, we performed an in depth qualitative and quantitative analysis of protein content and abundance in EDL, soleus and EOMs from WT and dHT mice. The results of the proteomic analysis reveal that, asides the drastic reduction in RyR1 content, profound changes occur in the content of many proteins particularly in fast-, slow-twitch and EOM muscles. Namely, we found that recessive *Ryr1* mutations lead to an increase content aspartyl-beta-hydroxylase (*Asph*), some ribosomal proteins and calmodulin kinase 2 delta. EDL and EOMs that are more severely affected and also shared changes in the content of other proteins, including collagens, heat shock proteins (BiP), FKBP12 and FKBP9 CamK2b as well as additional ribosomal proteins. We believe that the reduced RyR1 calcium channel content has a domino effect leading to changes in content of many other proteins, particularly in EDL and EOMs.

### Recessive *Ryr1* mutations affect the expression of collagen, chaperons and ribosomal proteins

“Reactome” interaction pathway analysis revealed that the major pathways affected by the presence of compound heterozygous *Ryr1* mutations in EDL muscles includes proteins involved in organization and degradation of the extracellular matrix (ECM) and indeed the content of collagen I, II, IV, V and XI was significantly reduced. The ECM plays an important role in muscle force transmission, maintenance and repair and collagen accounts for 1-10% muscle dry weight fibers, forming a highly ordered network surrounding individual muscle fibers and muscle bundles (38, 39). Exactly how defects in the collagen network impact muscle function is not clear, nevertheless patients bearing mutations in Collagen VI (*COL6A1, COL6A2* and *COL6A3*) suffer from Ulrich and Bethlem myopathies (40) and exhibit muscle contractures involving elbows and ankles, a clinical sign that has been also described in patients suffering of congenital myopathies linked to recessive *RYR1* mutations (15, 41). In addition, a frequent common feature of patients with congenital myopathies carrying recessive *RYR1* mutations is the appearance of a number of skeletal abnormalities at birth, including scoliosis and congenital dislocation of the hip, kyphosis, clubfoot, flattening of the arch of the foot (or an abnormally high arch of the foo). Patients also exhibit joint laxity that may lead to dislocation of the patella, or, more rarely abnormal tightening of certain joints, resulting in contractures especially of the Achilles tendon (42). Recent RNAseq and proteomic studies have shown that RyR1s are expressed in bone (www.proteomicsdb.org), a result which is consistent with the idea that RyRs mediated calcium signaling might be involved in bone remodeling (43, 44). We speculate that recessive *Ryr1* mutations might downregulate Col1alfa expression not only in skeletal muscle but also in cells involved in bone formation and /or remodeling. The decrease of Col1alfa expression in bone tissue in recessive *Ryr1* mutant mice, may cause skeleton defects similar to those described in a mouse model having a severe deficit in Col1alfa expression, which exhibit limb deformities, reduced body size, kyphosis and scoliosis (45).

Muscles from the dHT did not show upregulation of proteins related to ER stress such as PERK, IRE1a, ATF6 though the content of BiP as well as that of several heat shock proteins was significantly increased in EDL and EOM muscles. Heat shock proteins are molecular chaperones that participate in the safeguard of cell integrity, playing numerous functions including protection from heat insults, prevention of aggregation and facilitation of protein folding. There are different categories of heat shock proteins, including small HSPs (HSPB1-10) that are involved in protein folding, prevention of aggregation. In skeletal muscle these small proteins have been shown to be involved in the maintenance of the cytoskeletal network and contractile elements and play a role in myogenic differentiation. Large HSP are present in many subcellular locations including mitochondria, nucleus, sarcoplasmic reticulum and myoplasm where they facilitate protein folding and re-folding, facilitate protein transport into the SR and mitochondria and prevent aggregate formation. Interestingly, intensive resistance training increases heat shock protein levels in muscle (46) whereas aging is associated with a decrease in HSP70 response in muscles following muscle contraction (47, 48). Proteomic profiling has shown that muscle diseases including dysferlinopathies, myofibrillar myopathies, spinal muscular atrophy, Duchenne muscular dystrophy and others are associated with up-regulation of distinct HSP (for review see 49). These results together with our findings suggest that altered muscle function caused by genetic mutations are accompanied by adaptive cellular responses aimed at counterbalancing muscle damage and/or restoring proper function. Of note, in soleus muscles from the dHT mice, HSP are not up-regulated; this may be due to the fact that soleus muscles are less damaged/stressed or because the content of HSPs in soleus muscles is constitutively higher than in EDL muscles. In particular, the expression of HSP70 is 15-fold higher in WT soleus compared to WT EDL muscles. The high expression of HSP70 might thus protect slow twitch muscles from extensive damage linked to the expression of mutant RyR1s, an event which may ultimately account for the fiber type I predominance observed in patients with congenital myopathies linked to *RYR1* mutations.

An interesting observation of the present study is that the content of ribosomal proteins constituting the 40S and 60S subunits is significantly increased in the three muscle types from the dHT compared to WT. In skeletal muscle up-regulation of ribosomal proteins accompanies hypertrophy and training whereas ribosomal proteins decrease with age (50). Thus, our results indicate that the presence of *Ryr1* mutations evoke a global adaptive response aimed at (i) preserving the integrity of intracellular protein compartments and (ii) increasing muscle protein turnover.

### Stoichiometry of ECC molecular complex in health and diseased muscles

In this study we used spiked-in labelled peptides for isobaric TMT mass spectrometry measurements to quantify the major protein components of the EEC molecular complex in EDL, soleus and EOM from WT and dHT mice. We established the absolute content of low abundant ECC-molecular complex proteins, including RyR1, Cacna1s, Stim1, Orai1. The calculated values for RyR1 and Cacna1s that we obtained are of the same order of magnitude as those previously determined by equilibrium ligand binding (34–36), confirming the reliability of this approach. In addition, we also provide for the first time the absolute quantification of Stim1 and Ora1, two crucial proteins involved in Store Operated Calcium Entry (SOCE). Our results are interesting because of the widespread attention gained by SOCE in skeletal muscle, not only because mutations in *STIM1* and *ORAI1* are the underlying feature of several genetic diseases associated with muscle weakness (51, 52), but also because experimental evidence has shown that Stim1 and Orai1 play an important role in refilling intracellular calcium stores in fast and slow twitch muscles (37, 53, 54). Quantitative isobaric TMT mass spectrometry revealed for the first time important differences in the content of Stim1, Stim2 and Orai1 among different muscle types. We are confident of our results because the data relative to Stim1 content were validated by staining western blots of total muscle homogenates with Stim1 specific antibodies. Our data show that EOMs contain the highest levels of Stim1, and, in agreement with previous data by Cully et al. (37), we found no major differences in Stim1 content between fast and slow twitch muscles. The Stim1 to Orai1 ratio in EOMs is 34, a value approximately 1.4-fold higher compared to that of EDLs. This higher content of Stim1 and Orai1 supports the idea that SOCE is a robust component of calcium signaling in EOMs and may bring about a constant calcium entry necessary to replenish sarcoplasmic reticulum stores necessary to support the continuous fast muscle contraction unique to EOMs, compared to other striated muscles. A mind-boggling result emerging from the quantitative analysis of Stim1 and Orai1 is that slow twitch muscles such a soleus express a very small amount of Orai1 protein which could not be quantified by LC-MS. This raises the important question as to the nature of the molecular component(s) interacting with Stim1 in order to operate SOCE in slow twitch muscles. At this point in time, we cannot exclude the possibility that in slow twitch muscles, Stim1 interacts with a molecular partner different from Orai1, or that SOCE might be operated by an Orai1 variant having a much higher divalent cation conductance compared to the “classical” Orai1 isoform expressed in EDL and EOM. Nevertheless, such a question is beyond the scope of the present investigation and cannot be answered by the data presented here.

*STAC3* mutations have been linked to Native American Myopathy (NAM), a severe congenital myopathy resulting in muscle weakness and skeleton alteration (55). Such mutations cause a decrease of the interaction between Stac3 with Cacna1s resulting in a functional deficit of EC coupling (56). On the basis the quantitative data we obtained using the LC-MS standard curve generated by spiked-in peptides, the Stac3 to Cacna1s stoichiometry ratio is 1.11, 1.22 and 1.67 in EDL, soleus and EOM respectively, and no differences were observed between WT and dHT mice. Stac3 interacts via its SH3 domain with a Kd ranging between 2-10 μM, with a binding site within the cytosolic II-III loop of the Cacna1s (57–58). Here we show that the molar content of Stac3 in EDL, soleus and EOMs is between 2.5 to 10-fold lower than its Kd for the Cacna1s II-III loop binding site (56, 57). Thus, the fractional occupancy of the Cacna1s binding site by Stac3 is lower than 50%, a value which is still sufficient to support normal EC coupling. Nevertheless, the extent of the fractional occupancy depends on the fiber type. In particular, if the Kd of the Cacna1s binding site for Stac3 is identical in EDL, soleus and EOM, then the fractional occupancy of the Cacna1s binding site for Stac3 in soleus and EOMs is lower than of EDL muscles, because the molar content of Stac3 in soleus and EOMs is three-fold lower compared to that of EDL (Table 5). *STAC3* mutations linked to NAM decrease the Kd of the SH3 domain of Stac3 for the cytosolic II-III loop of Cacna1s (57) further lowering the fractional occupancy of Stac3 binding site of Cacna1s to a low level close to zero, a condition that would disrupt EC coupling in NAM patients (55).

Multiplexed proteomic analysis is a powerful approach for the quantitative proteomic analysis of a variety of biological samples. In particular, absolute quantification can be achieved by measuring the content of a protein relative to a spiked-in peptide with known absolute concentration. A limitation of the multiplexed isobaric mass tag-based protein quantification is the reliable detection of very low abundant proteins, such as transcriptional factors and other molecules involved in cellular signaling. Because of this intrinsic hurdle of multiplex isobaric mass tag spectrometry, in this study we missed nuclear proteins in addition to protein components of signaling pathways. An additional drawback of this study is that it gives a static image of muscle protein content in young adult mice without conveying information about the dynamics of protein changes or changes in post-translational modifications occurring during muscle disease.

In conclusion our quantitative proteomic study: 1) shows that recessive *Ryr1* mutations not only decrease the content of RyR1 protein in muscle, but also affect the content of many other proteins; 2) provides insight as to the potential pathological mechanism of congenital myopathies linked to mutation of other components of the ECC machinery.

## MATERIALS AND METHODS

### Compliance with Ethical standards

All experiments involving animals were carried out on 12 weeks old male wild type and dHT mice littermates. Experimental procedures were approved by the Cantonal Veterinary Authority of Basel Stadt (BS Kantonales Veterinäramt Permit numbers 1728). All experiments were performed in accordance with relevant guidelines and regulations.

### Proteomics analysis using tandem mass tags

EDL, soleus and EOM muscles from 5 male WT and 5 male dHT, 12 weeks old mice were excised, weighed, snap frozen in liquid nitrogen and mechanically grinded. Approximately 10 mg of EDL, 8 mg for of Soleus and 6 mg of EOM muscle tissue was grinded and subsequently lysed in 200 μl of lysis buffer containing 100 mM TRIS, 1% sodium deoxycholate (SDC), 10 mM TCEP and 15 mM chloroacetamide, followed by sonication (Bioruptor, 20 cycles, 30 seconds on/off, Diagenode, Belgium) and heating to 95°C for 10 minutes. After cooling, protein samples were digested by incubated overnight at 37°C with sequencing-grade modified trypsin (1/50, w/w; Promega, Madison, Wisconsin). Samples were acidified using 5% TFA and peptides cleaned up using the Phoenix 96× kit (PreOmics, Martinsried, Germany) following the manufacturer’s instructions. After drying the peptides in a SpeedVac, samples were stored at −80°C.

Dried peptides were dissolved in 100 μl of 0.1% formic acid and the peptide concentration determined by UV-nanodrop analysis. Sample aliquots containing 25 μg of peptides were dried and labeled with tandem mass isobaric tags (TMT 10-plex, Thermo Fisher Scientific) according to the manufacturer’s instructions. To control for ratio distortion during quantification, a peptide calibration mixture consisting of six digested standard proteins mixed in different amounts were added to each sample before TMT labeling as recently described (18). After pooling the differentially TMT labeled peptide samples, peptides were again desalted on C18 reversed-phase spin columns according to the manufacturer’s instructions (Macrospin, Harvard Apparatus) and dried under vacuum. Half of the pooled TMT-labeled peptides (125 μg of peptides) were fractionated by high-pH reversed phase separation using a XBridge Peptide BEH C18 column (3,5 μm, 130 Å, 1 mm × 150 mm, Waters) on an Agilent 1260 Infinity HPLC system. 125 ug of peptides were loaded onto the column in buffer A (ammonium formate (20 mM, pH 10, in water) and eluted using a two-step linear gradient starting from 2% to 10% in 5 minutes and then to 50% (v/v) buffer B (90% acetonitrile / 10% ammonium formate (20 mM, pH 10) over 55 minutes at a flow rate of 42 μl/min. Elution of peptides was monitored with a UV detector (215 nm, 254 nm). A total of 36 fractions were collected, pooled into 12 fractions using a post-concatenation strategy as previously described (59) and dried under vacuum.

The generated 12 peptide samples fractions were analyzed by LC-MS as described previously (18). Chromatographic separation of peptides was carried out using an EASY nano-LC 1000 system (Thermo Fisher Scientific), equipped with a heated RP-HPLC column (75 μm × 37 cm) packed in-house with 1.9 μm C18 resin (Reprosil-AQ Pur, Dr. Maisch). Aliquots of 1 μg of total peptides of each fraction were analyzed per LC-MS/MS run using a linear gradient ranging from 95% solvent A (0.15% formic acid, 2% acetonitrile) and 5% solvent B (98% acetonitrile, 2% water, 0.15% formic acid) to 30% solvent B over 90 minutes at a flow rate of 200 nl/min. Mass spectrometry analysis was performed on Q-Exactive HF mass spectrometer equipped with a nanoelectrospray ion source (both Thermo Fisher Scientific). Each MS1 scan was followed by high-collision-dissociation (HCD) of the 10 most abundant precursor ions with dynamic exclusion for 20 seconds. Total cycle time was approximately 1 s. For MS1, 3e6 ions were accumulated in the Orbitrap cell over a maximum time of 100 ms and scanned at a resolution of 120,000 FWHM (at 200 m/z). MS2 scans were acquired at a target setting of 1e5 ions, accumulation time of 100 ms and a resolution of 30,000 FWHM (at 200 m/z). Singly charged ions and ions with unassigned charge state were excluded from triggering MS2 events. The normalized collision energy was set to 35%, the mass isolation window was set to 1.1 m/z and one microscan was acquired for each spectrum.

The acquired raw-files were searched against a protein database containing sequences of the predicted SwissProt entries of mus musculus (www.ebi.ac.uk, release date 2019/03/27), Myh2 and Myh13 from Trembl, the six calibration mix proteins (18) and commonly observed contaminants (in total 17,414 sequences) using the SpectroMine software (Biognosys, version 1.0.20235.13.16424) and the TMT 10-plex default settings. In brief, the precursor ion tolerance was set to 10 ppm and fragment ion tolerance was set to 0.02 Da. The search criteria were set as follows: full tryptic specificity was required (cleavage after lysine or arginine residues unless followed by proline), 3 missed cleavages were allowed, carbamidomethylation (C), TMT6plex (K and peptide n-terminus) were set as fixed modification and oxidation (M) as a variable modification. The false identification rate was set to 1% by the software based on the number of decoy hits. Proteins that contained similar peptides and could not be differentiated based on MS/MS analysis alone were grouped to satisfy the principles of parsimony. Proteins sharing significant peptide evidence were grouped into clusters. Acquired reporter ion intensities in the experiments were employed for automated quantification and statistically analyzed using a modified version of our in-house developed SafeQuant R script (v2.3)(18). This analysis included adjustment of reporter ion intensities, global data normalization by equalizing the total reporter ion intensity across all channels, summation of reporter ion intensities per protein and channel, calculation of protein abundance ratios and testing for differential abundance using empirical Bayes moderated t-statistics. Finally, the calculated p-values were corrected for multiple testing using the Benjamini-Hochberg method.

### Targeted PRM-LC-MS analysis of Ryr1 and Cacna1s, Stim1 and Orai1

In a first step, parallel reaction-monitoring (PRM) assays (60) were generated from a mixture containing 50 fmol of each proteotypic heavy reference peptide of the target proteins (AIWAEYDPEAK, GEGIPTTAK, TGGLFGQVDNFLER (for Cacna1s); AGDVQSGGSDQER, GPHLVGPSR, SNQDLITENLLPGR, TLLWTFIK, VVAEEEQLR (for Ryr1); LISVEDLWK, AIDTVIFGPPIITR, ITEPQIGIGSQR, LSFEAVR, YAEEEIEQVR (for Stim1); QFQELNELAEFAR, IQDQIDHR, SLVSHK (for Orai1); JPT Peptide Technologies GmbH) plus iRT peptides (Biognosys, Schlieren, Switzerland). Peptides were subjected to LC–MS/MS analysis using a Q Exactive Plus mass spectrometer fitted with an EASY-nLC 1000 (both Thermo Fisher Scientific) and a custom-made column heater set to 60°C. Peptides were resolved using a RP-HPLC column (75μm × 30cm) packed in-house with C18 resin (ReproSil-Pur C18–AQ, 1.9 μm resin; Dr. Maisch GmbH) at a flow rate of 0.2 μLmin-1. A linear gradient ranging from 5% buffer B to 45% buffer B over 60 minutes was used for peptide separation. Buffer A was 0.1% formic acid in water and buffer B was 80% acetonitrile, 0.1% formic acid in water. The mass spectrometer was operated in DDA mode with a total cycle time of approximately 1 s. Each MS1 scan was followed by high-collision-dissociation (HCD) of the 20 most abundant precursor ions with dynamic exclusion set to 5 seconds. For MS1, 3e6 ions were accumulated in the Orbitrap over a maximum time of 254 ms and scanned at a resolution of 70,000 FWHM (at 200 m/z). MS2 scans were acquired at a target setting of 1e5 ions, maximum accumulation time of 110 ms and a resolution of 35,000 FWHM (at 200 m/z). Singly charged ions, ions with charge state ≥ 6 and ions with unassigned charge state were excluded from triggering MS2 events. The normalized collision energy was set to 27%, the mass isolation window was set to 1.4 m/z and one microscan was acquired for each spectrum. The acquired raw-files were searched using the MaxQuant software (Version 1.6.2.3) against the same protein sequence database as decribed above using default parameters except protein, peptide and site FDR were set to 1 and Lys8 and Arg10 were added as variable modifications. The best 6 transitions for each peptide were selected automatically using an in-house software tool and imported into SpectroDive (version 8, Biognosys, Schlieren). A scheduled (window width 12 min) mass isolation list containing the iRT peptides was exported form SpectroDive and imported into the Q Exactive plus operating software for PRM analysis.

Peptide samples for PRM analysis were resuspended in 0.1% aqueous formic acid, spiked with iRT peptides and the heavy reference peptide mix at a concentration of 10 fmol of heavy reference peptides per 1 μg of total endogenous peptide mass and subjected to LC–MS/MS analysis on the same LC-MS system described above using the following settings: The resolution of the orbitrap was set to 140,000 FWHM (at 200 m/z), the fill time was set to 500 ms to reach an AGC target of 3e6, the normalized collision energy was set to 27%, ion isolation window was set to 0.4 m/z and the first mass was fixed to 100 m/z. A MS1 scan at 35,000 resolution (FWHM at 200 m/z), AGC target 3e6 and fill time of 50 ms was included in each MS cycle. All raw-files were imported into SpectroDive for protein / peptide quantification. To control for variation in injected sample amounts, the total ion chromatogram (only comprising ions with two to five charges) of each sample was determined and used for normalization. To this end, the generated raw files were imported into the Progenesis QI software (Nonlinear Dynamics (Waters), Version 2.0), the intensity of all precursor ions with a charge of +2 to +5 were extracted, summed for each sample and used for normalization. Normalized ratios were transformed from the linear to the log-scale, normalized relative to the control condition and the median ratio among peptides corresponding to one protein was used for protein quantification.

### Western blot analysis of Stim1 and Stim1L

Total homogenates of EDL, soleus and EOM muscles from WT mice were prepared in cracking buffer as previously described (16, 17). Proteins were separated on a 7.5% SDS-PAG, blotted onto nitrocellulose and probed with an antibody recognizing Stim1 and Stim1L (1/2000 anti-STIM1, Millipore, #AB9870), followed by incubation with an anti-rabbit IgG HRP-linked antibody (1/6000, Cell Signaling Technology, #7074). Bands were visualized by chemiluminescence. Blots were subsequently stripped and probed with anti-MyHC (all) (1/5000, DSHB, #MF20) for loading normalization as previously described (16, 17). Statistical analysis was performed using a one-way ANOVA test.

### Data-analyses

Matlab 2021b (Mathworks) (61) was used to process the proteomics data and to generate heatmap, volcano plots and Venn diagrams.

## List of Abbreviations

CM: congenital myopathies
dHT: compound heterozygous double *Ryr1* mutant mice
ECC: excitation-contraction coupling
ECM: extracellular matrix
EDL: *extensor digitorum longus*
EOM: extraocular muscles
FDB: *flexor digitorum brevis*
MS: mass spectrometry
MmD: multiminicore disease
MyHC: myosin heavy chain
NAM: Native American Myopathy
RyR1: ryanodine receptor 1
WT: wild type mice

## Author contributions

Conceptualization: FZ and ST

Methodology: JE, AR, SK, MF, AS, ST, FZ

Investigation: AR, ST, SB, MF, LP, SB, CB, FN, FP, FZ

Funding acquisition: ST, FZ

Project administration: ST, FZ

Supervision: ST, FZ, MF

Writing – original draft: FZ, ST

Writing – review & editing: AR, ST, SB, MF, LP, SB, CB, FN, FP, FZ

## Acknowledgements

The authors wish to acknowledge Dr. Eric Ahrné for constructive discussions.

## Data and materials availability

All data, code, and materials used in the analysis are available in some form to any researcher for purposes of reproducing or extending the analysis. There are no restrictions on materials, such as materials transfer agreements (MTAs). The Mass Spectrometry proteomic data have been deposited to the ProteomeXchange Consortium via the Pride partner repository (http://ebi.ac.uk/pride) with the dataset identifier PXD036789 and 10.6019/PXD036789.

## Supplementary Material

**Supplementary Figure 1:**
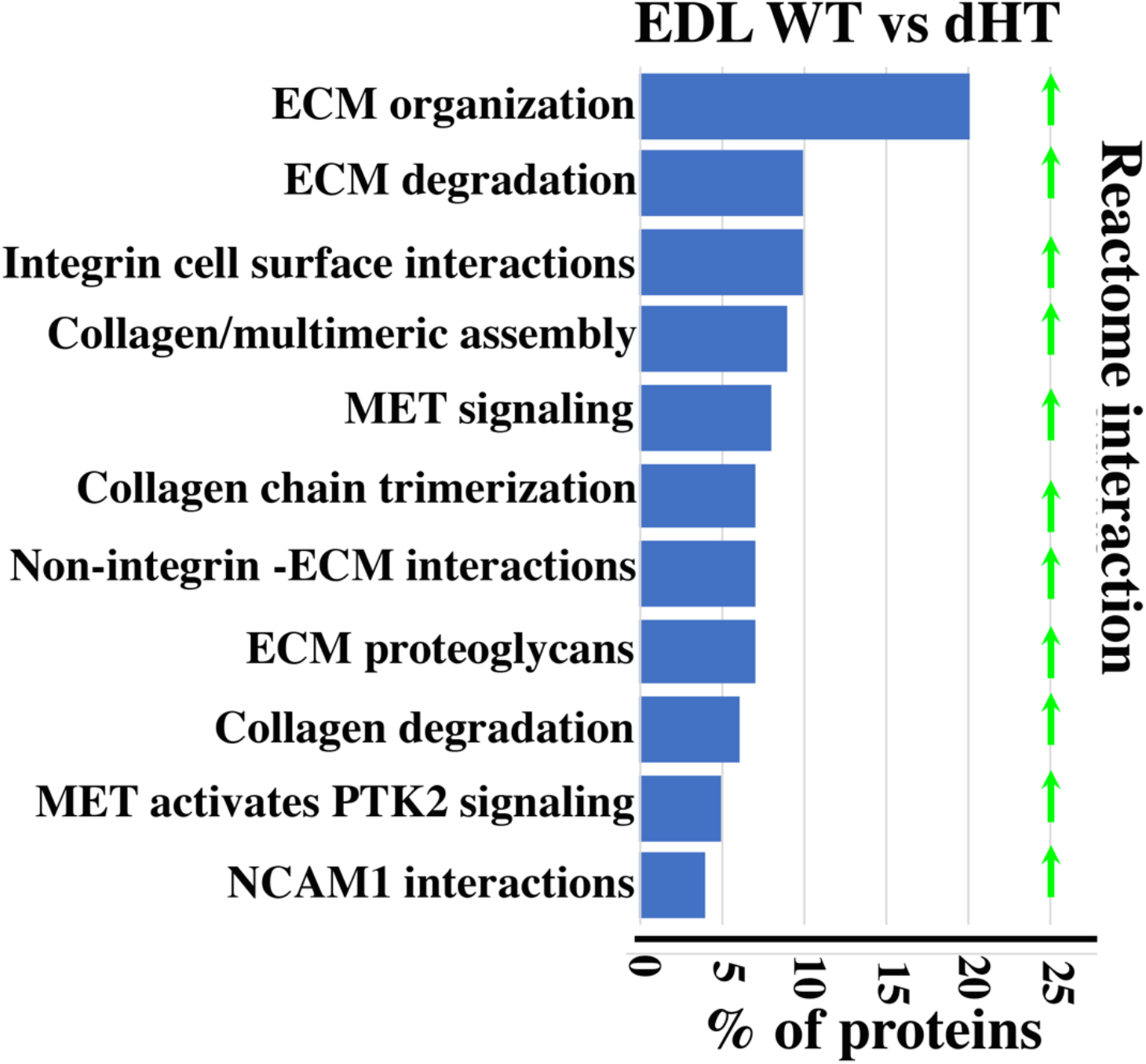
Reactome pathway analysis showing major pathways which differ between EDL muscles in WT versus dHT mice. A q-value of equal or less than 0.05 was used to filter significant changes prior to the pathway analyses

**Supplementary Figure 2:**
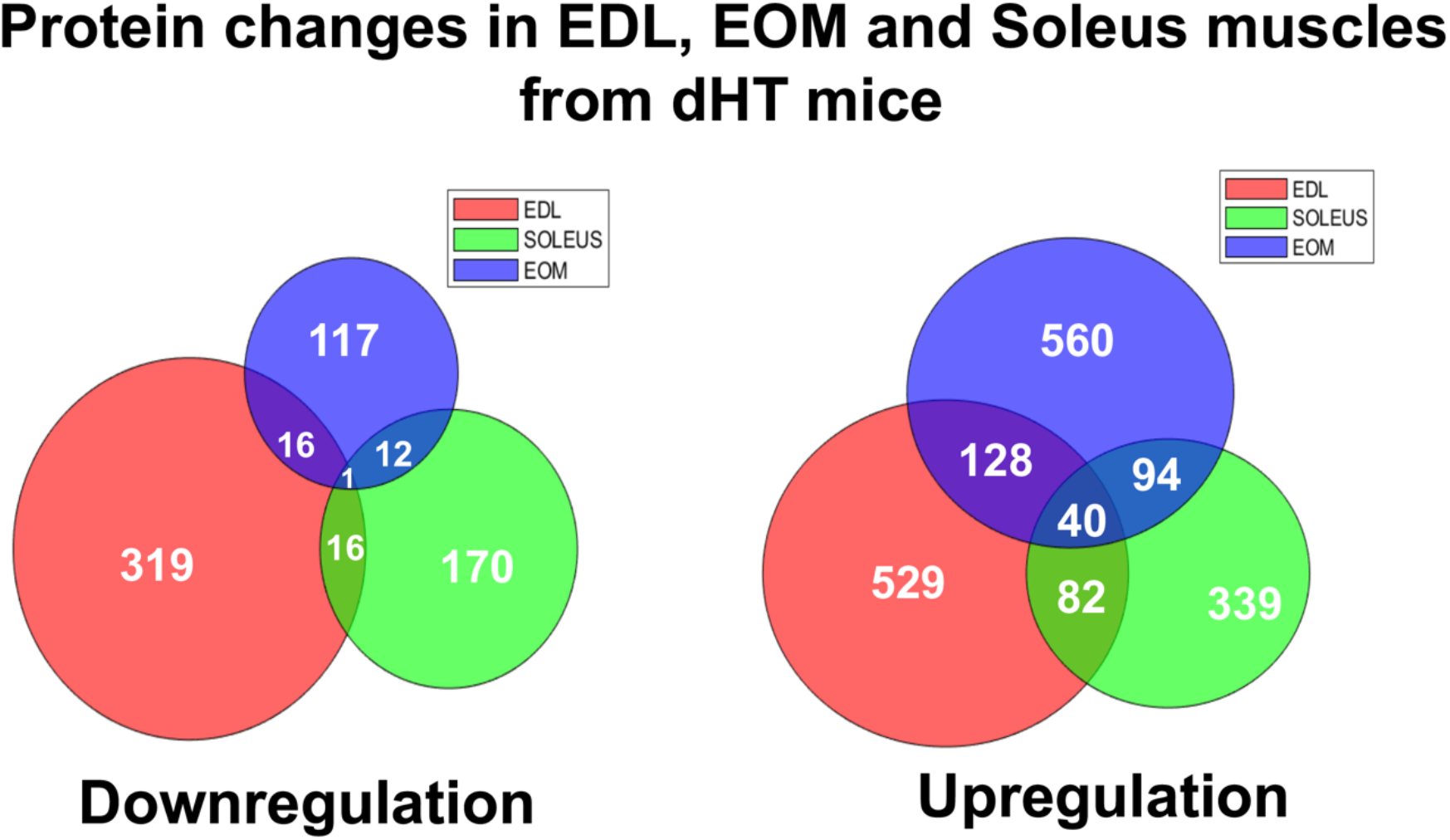
Venn diagrams showing the total number of proteins exhibiting a significant change in content in muscles from dHT mice compared to WT littermates. There is only one down-regulated protein common to EDL, soleus and EOM muscles, namely RyR1, whereas the content of 40 proteins was up-regulated in all three muscle types.

**Supplementary Figure 3:**
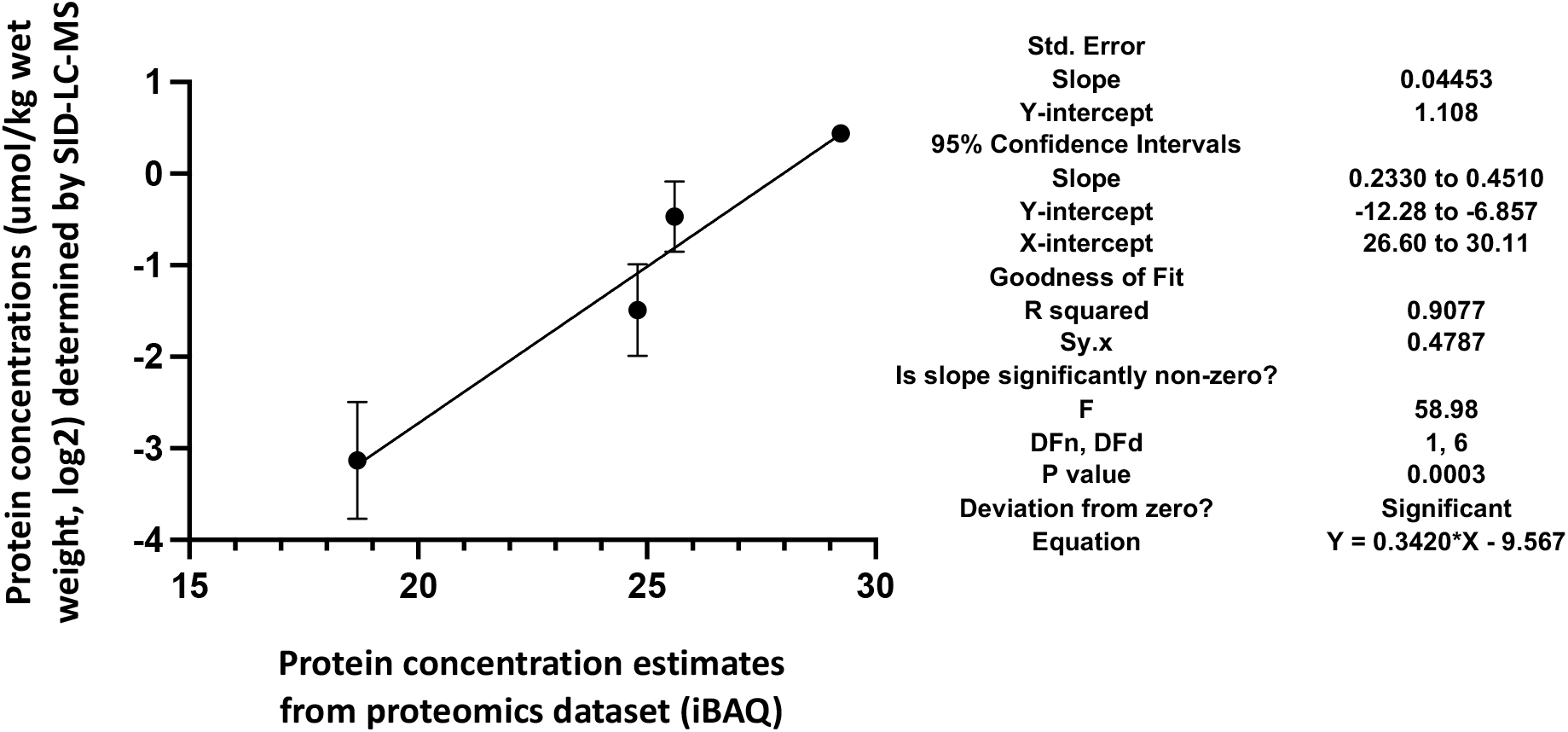
Correlation of the actual cellular abundances of 4 selected proteins (in μmol/kg wet weight) determined by PRM/SID (n=2) and the iBAQ values (n=5) determined by label-free/TMT quantification (both in logarithmic scale, base 2) from the global proteomics discovery dataset for EDL samples. Error bars are indicated for the y-axis, but for the x-axis, due to their low scale (range from 0.058-0.086), they are not shown by the software (PRISM, GraphPad Software, v9). The simple linear regression results obtained by PRISM (GraphPad Software, v9) are shown on the right.

